# Multi-trait modeling and machine learning discover new markers associated with stem traits in alfalfa

**DOI:** 10.1101/2024.05.03.592319

**Authors:** Cesar A. Medina, Deborah J. Heuschele, Dongyan Zhao, Meng Lin, Craig T. Beil, Moira J. Sheehan, Zhanyou Xu

## Abstract

Alfalfa biomass can be fractionated into leaf and stem components. Leaves comprise a protein-rich and highly digestible portion of biomass for ruminant animals, while stems constitute a high fiber and less digestible fraction, representing 50 to 70% of the biomass. However, little attention has focused on stem-related traits, which are a key aspect in improving the nutritional value and intake potential of alfalfa. This study aimed to identify molecular markers associated with four morphological traits in a panel of five populations of alfalfa generated over two cycles of divergent selection based on 16-h and 96-h in vitro neutral detergent fiber digestibility in stems. Phenotypic traits of stem color, presence of stem pith cells, winter standability, and winter injury were modeled using univariate and multivariate spatial mixed linear models (MLM), and the predicted values were used as response variables in genome-wide association studies (GWAS). The alfalfa panel was genotyped using a 3K DArTag SNP markers for the evaluation of the genetic structure and GWAS. Principal component and population structure analyses revealed differentiations between populations selected for high- and low-digestibility. Thirteen molecular markers were significantly associated with stem traits using either univariate or multivariate MLM. Additionally, support vector machine (SVM) and random forest (RF) algorithms were implemented to determine marker importance scores for stem traits and validate the GWAS results. The top-ranked markers from SVM and RF aligned with GWAS findings for solid stem pith, winter standability, and winter injury. Additionally, SVM identified additional markers with high variable importance for solid stem pith and winter injury. Most molecular markers were located in coding regions. These markers can facilitate marker-assisted selection to expedite breeding programs to increase winter hardiness or stem palatability.

**Author Summary:** The alfalfa stem constitutes a significant portion of forage yield, accounting for 50 to 70% of biomass yield, and influences various traits including plant height, standability, and digestibility. In our study, we identified thirteen molecular markers associated with stem color, presence of stem pith parenchyma, winter standability, and winter injury in a panel of five divergent stem digestibility populations. Multivariate trait modeling enhances the correlation among traits, thereby expanding the number of markers associated via GWAS. Similarly, machine learning algorithms increase the confidence of markers initially identified by GWAS and uncover new candidate regions that could serve as associated markers. Genes harboring markers associated to the four stem traits play roles in plant growth, response to plant injury, or tolerance to cold, underscoring their potential utility in enhancing traits such as cold tolerance and forage quality in alfalfa.

## Introduction

Alfalfa (*Medicago sativa* L.) is the third-largest field crop produced in the United States. It is grown for hay, silage, and pasture due to its high nutritive value for milk and muscle mass production in livestock [1]. During harvest, all aerial parts of the plant are collected. Alfalfa biomass yield can be fractionated into leaf and stem components. The leaves comprise a protein-rich, highly digestible portion for ruminant animals, while the stem component represents 50 to 70% of the biomass and comprises a high-fiber, less digestible fraction [2]. Therefore, improving stem digestibility will increase palatability, dry matter, and available energy of the plant especially at later maturity stages.

There are different approaches to increase forage digestibility, including gene knockdowns in the lignin biosynthetic pathway [3,4], increasing the proportion of non-lignified tissues [5], increasing the leaf/stem ratio [6], or reducing lignin concentration to increase fiber digestibility using traditional breeding [7]. Previously, a panel of five populations was developed to enhance stem digestibility using recurrent selection for in vitro neutral detergent fiber digestibility (IVNDFD) in alfalfa stems [5,8]. Jung & Lamb (2006) generated a cycle zero population (UMN3097) by mixing seeds from six commercial alfalfa varieties. Genotypes of UMN3097 were categorize into four groups: low rapid (16-h) IVNDFD, high rapid (16-h) IVNDFD, low potential (9-h) IVNDFD, and high potential (96-h) IVNDFD. Two cycle 1 divergent stem IVNDFD populations were developed through intercrossing plants with high 16-h and 96-h digestibility (H×H) (UMN3355) and low 16-h and low 96-h digestibility (L×L) (UMN3358). Two cycle 2 divergent stem IVNDFD populations were generated by intermating H×H genotypes of UMN3355 and L×L genotypes of UMN3358 (UMN4016 and UMN4019, respectively). This approach aimed to increase biomass yields while maintaining forage quality and enhancing seasonal stability in stem digestibility.

The stem structure of alfalfa changes during development from young to mature tissues. The young stem has a square shape in cross-section, with major vascular bundles located in the angles, while the mature stem tissues are rounded due to cambial growth [9]. The center of the stem is occupied by the pith, composed of large, compactly arranged parenchyma cells. The parenchyma cells in the pith are unlignified and as stems mature these cells may die, leading to the formation of air-filled cavities in the stem [10]. The generation of a hollow stem is related to an increased stem lodging and stalk-rot in maize [11]. In alfalfa, stem pith parenchyma is an important trait related to stem maturity, stem water and nutrient content, and palatability [10].

Modifications of tissue composition can affect other traits like plant size or susceptibility to diseases [12]. Therefore, it is necessary to evaluate different traits that could affect the performance of the selected populations. In the Northern Great Plains of the United States, the ability to develop and maintain an adequate level of freezing resistance is imperative to withstand stressful mid-winter temperatures. During the fall, alfalfa increases tolerance to low temperatures and prepares to enable roots and crowns to survive temperatures as low as -20°C. Winter injury occurs due to cold temperatures in winter that cause freezing of root cells or crown buds, resulting in subsequent damage due to freezing. Sublethal winter injury can decrease vigor during the subsequent growing season [13].

The availability of the Diversity Array Technology (DArTag) genotyping platform for alfalfa enables the acquisition of highly consistent 3,000 SNPs to implement genome-wide association studies (GWAS) or genomic selection [14]. GWAS can identify marker-trait associations using mixed linear models (MLM) that includes population structure and a kinship matrix to correct false associations [15]. However, GWAS only analyzes the linear relations for each SNP individually and cannot detect SNP-SNP interactions or small-effect variants. On the other hand, machine learning (ML) models have fewer assumptions about the normality of data and can capture non-linear interactions between the predictor variables (i.e. SNPs) and the response variable. Additionally, ML can capture the combined minor effects of multiple genetic markers, allowing for the development of multi-locus methodologies that consider all SNPs in the model to estimate the importance scores of SNPs. Support vector machines (SVM) and random forest (RF) are two of the most effective machine learning models for predictive analytics capable of improving GWAS results [16,17].

The objective of this study was to identify molecular markers associated with four morphological traits (stem color, presence of stem pith parenchyma [hereafter referred to as stem fill], winter standability, and winter injury) in a panel of five populations of alfalfa generated over two cycles of divergent selection for 16-h and 96-h IVNDFD. In this work, we compared the genotypic response of univariate and multivariate spatial MLM and the markers associated with GWAS and ML models.

## Results

### Genotype information

In this study, we genotyped 1,502 individuals from five divergent stem digestibility populations using a mid-density DArTag platform. The five populations used were the cycle zero population (C0); the cycle 1 population with high16-h and 96-h IVNDFD (C1 H×H); the cycle 1 population with low 16-h and 96-h IVNDFD (C1 L×L); the cycle 2 population with high 16-h and 96-h IVNDFD (C2 H×H); and the cycle 2 population with low 16-h and 96-h IVNDFD (C2 L×L) (see plant materials and field experiment in materials and methods section).

Initially, the Allele Match Count Collapsed (AMCC) file contained 3,000 target SNP markers; however, after minor allele frequency (MAF) and collinearity filtering, the genotypic matrix kept 2,434 SNP markers (81%). Markers were not evenly distributed along or among chromosomes (Figure 1a). Chromosome 6 had the lowest number of markers (153), with a density of 1.91 SNP/Mb, and a maximum gap between two markers of 4,339 kb around the centromeric region. Chromosome 4 had the highest number of markers (362), with a density of 4.01 SNP/Mb, and a maximum gap between two markers of 1,212 kb. Allele dosage in the five alfalfa populations showed an increase in the heterozygous markers in the C1 and C2 L×L populations (Figure 1b). This was corroborated by heterozygosity-based statistics. Observed heterozygosity (H_O_) was greater than the expected heterozygosity within the population (H_S_) in all populations, which is often observed in highly diverse populations. H_O_ was the lowest in the UMN3097 (C0) population (0.391) and highest in the UMN3358 (C1 L×L) population (H_O_ = 0.441). Additionally, the UMN3358 population had the highest value of genetic diversity (H_S_ = 0.387) (Figure 1c).

**Figure 1.**
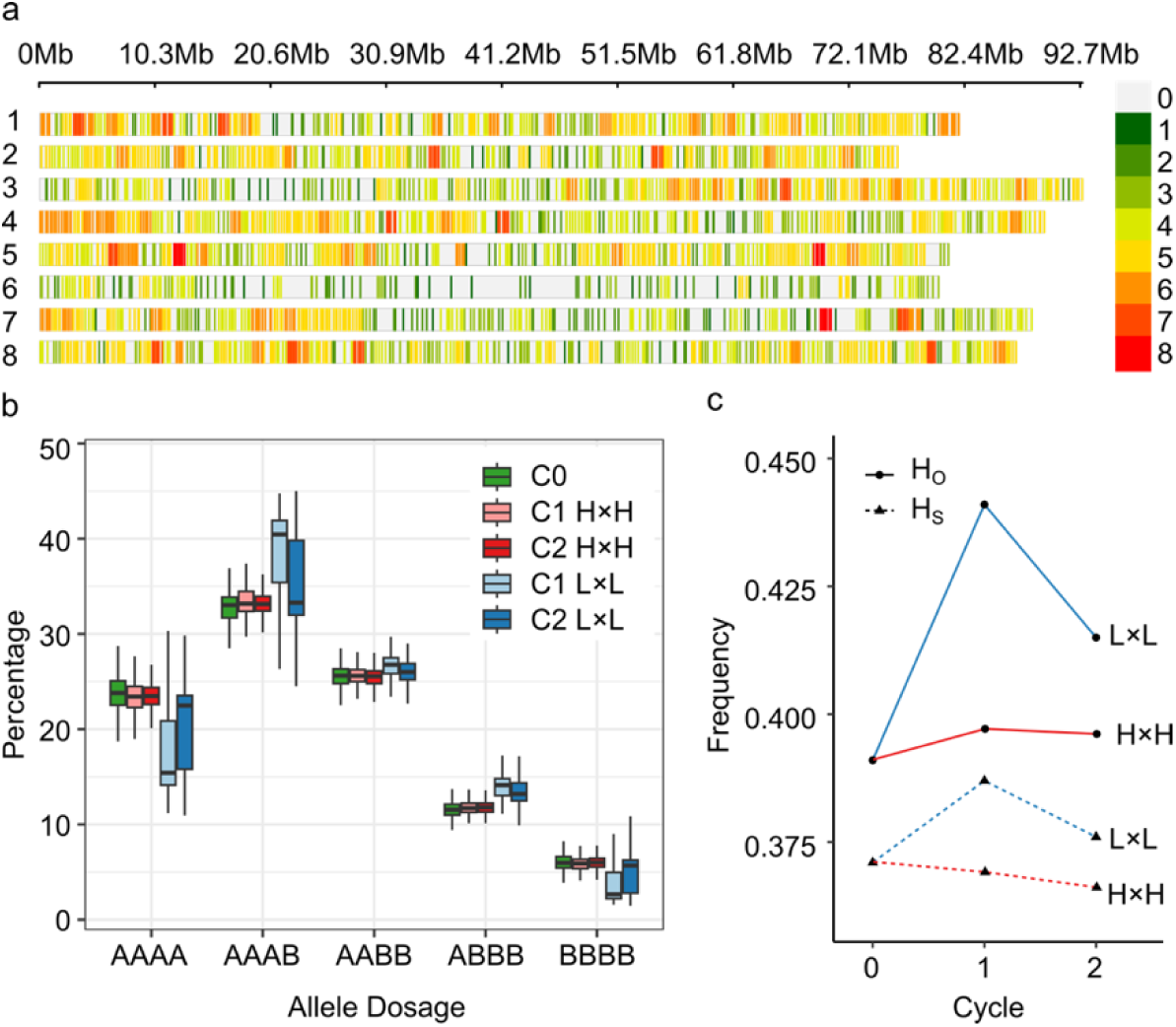
Genotypic information of SNP markers in divergent stem digestibility populations. **a.** SNP-density plot of 2,434 biallelic SNPs the eight chromosomes of *Medicago sativa*; colors represent number of SNPs within a 1 megabase (Mb) window size. Loci with high-density SNPs are shown in red, and loci with low-density SNPs are shown in green. **b.** Boxplot of allele dosage in five alfalfa populations. A and B represents the reference and alternative allele, respectively. **c.** Observed (H_O_) and expected heterozygosity (H_S_) by selection cycle. C0, cycle zero population (UMN3097); C1 H×H, cycle 1 population with high 16-h and 96-h vitro neutral detergent fiber digestibility (IVNDFD) (UMN3355); C2 H×H, cycle 2 population with high 16-h and 96-h IVNDFD (UMN4016); C1 L×L, cycle 1 population with low 16-h and 96-h IVNDFD (UMN3358); and C2 L×L, cycle 2 population with low 16-h and 96-h IVNDFD (UMN4019).

### Population structure

A Principal Component Analysis (PCA) was conducted with 1,502 genotypes to define the genetic relationships among populations. The first and second components contributed 6.29% of the total genetic variance. The individuals of the five populations clustered according to cycle and selection criteria (Figure 2a). C0 genotypes were grouped in the middle of the scatter plot (coordinates 0, 0), individuals of cycles 1 and 2 with high digestibility were grouped in the right section of the scatter plot, and individuals of cycles 1 and 2 with low digestibility were grouped in the left section of the scatter plot. C2 populations were more divergent from the base population because of selection. According to Evanno’s ΔK method, the most likely value of K was two, indicating that the 1,502 genotypes could be grouped into two clusters based on differences in their markers (Supplementary Figure 1). All genotypes were grouped by population and sorted according to the membership to the two clusters.

**Figure 2.**
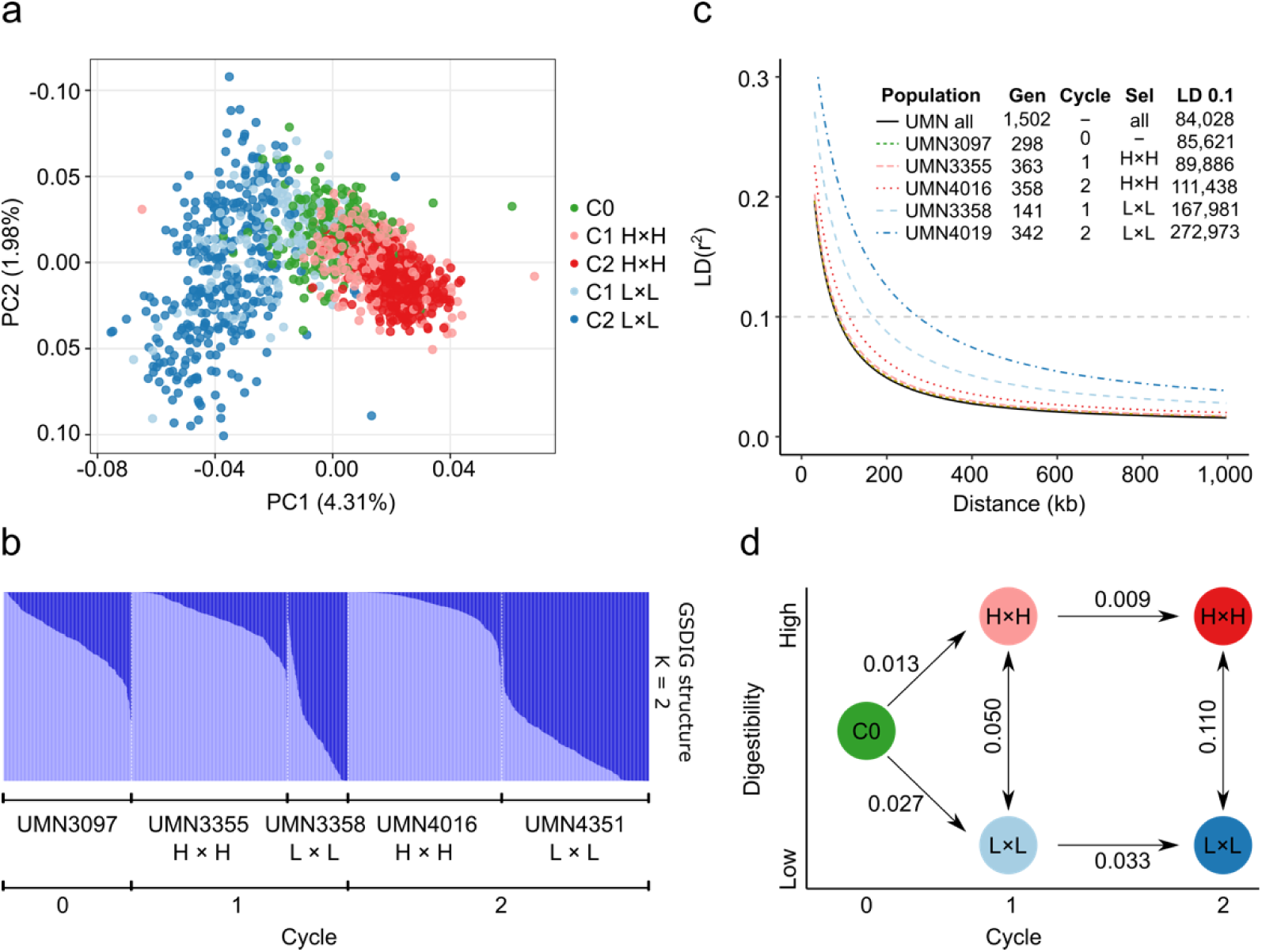
Population structure analysis of divergent stem digestibility populations. **a.** Principal component analysis for the panel composed of 1,502 genotypes. Colors correspond to the five alfalfa populations. **b.** Population structure bar plot at K = 2 inferred by the STRUCTURE program for 1,502 genotypes. Each bar represents one genotype, and the bar colors represent the likelihood of membership in each subpopulation. **c.** Linkage disequilibrium decay of combined (All) populations for five alfalfa populations. The values of half-decay and distances in bp at r^2^ = 0.1 are shown in the inner table. The half values were estimated based on the maximum values of LD decay. Gen = Number of genotypes. Sel = selection criteria. **d.** Graph of Rho interpopulation distance.

Linkage disequilibrium (LD) was determined by fitting a non-linear model between the square of the correlation coefficient among pairs of SNPs and physical distances on the *M. sativa* genome. LD decay was measured in all genotypes and in each of the five individual populations. LD decay at r^2^ = 0.1 among all genotypes was the lowest (84 kb), and the values increased with selection cycle. LD decay in the C0 population was the second lowest (86 kb), while C1 and C2 L×L populations had the highest LD decay (168 and 273 kb, respectively) (Figure 2c). LD decay was notably higher in L×L populations, doubling and tripling the LD block size in C1 and C2, respectively. Similarly, genetic differentiation values (Rho) were highest between L×L and H×H in C2 (0.11) and lowest between C1 and C2 in H×H populations (0.009). Rho values were doubled when comparing L×L and H×H populations in C1 vs C2, and Rho values were greater in the generation of L×L populations (Figure 2d).

### Genotypic modeling

The best linear unbiased estimates (BLUEs) values were obtained using a single trait (ST) or univariate spatial mixed linear model (MLM). The genotypes and replicates were defined as fixed components, while the nugget effect, columns, and rows were considered random components, and a spatial autoregressive correlation matrix was included as a residual component. Pairwise comparisons among populations showed a significant difference between high and low-digestibility C2 populations in stem color. High and low digestibility C2 populations were significantly different (p-value <0.05) in stem color; and in winter standability (Supplementary Figure 2). The Wald test for fixed effects identified that genotypic variation was highly significant (p-value <0.001) for stem color, stem fill, and winter injury, but not for winter standability (Supplementary Table 1). The nugget effect was included in the spatial MLM and corresponded to the measure of error variance in the spatial modeling at an infinitesimal separation distance between plots. A log likelihood ratio test showed a significant improvement (p-value <0.001) to the model fit including the nugget effect. Genetic variance was the highest for winter injury (Vg = 0.54) and the lowest for winter standability (Vg = 7.37 ×10^-8^) (Figure 3a). The broad sense heritability (H^2^) was high for winter injury (0.78) indicating most of the phenotypic variation of this trait was genetically controlled. H^2^ was medium for stem color (0.22) and stem fill (0.47), and the lowest for winter standability (0.04) (Figure 3b).

**Figure 3.**
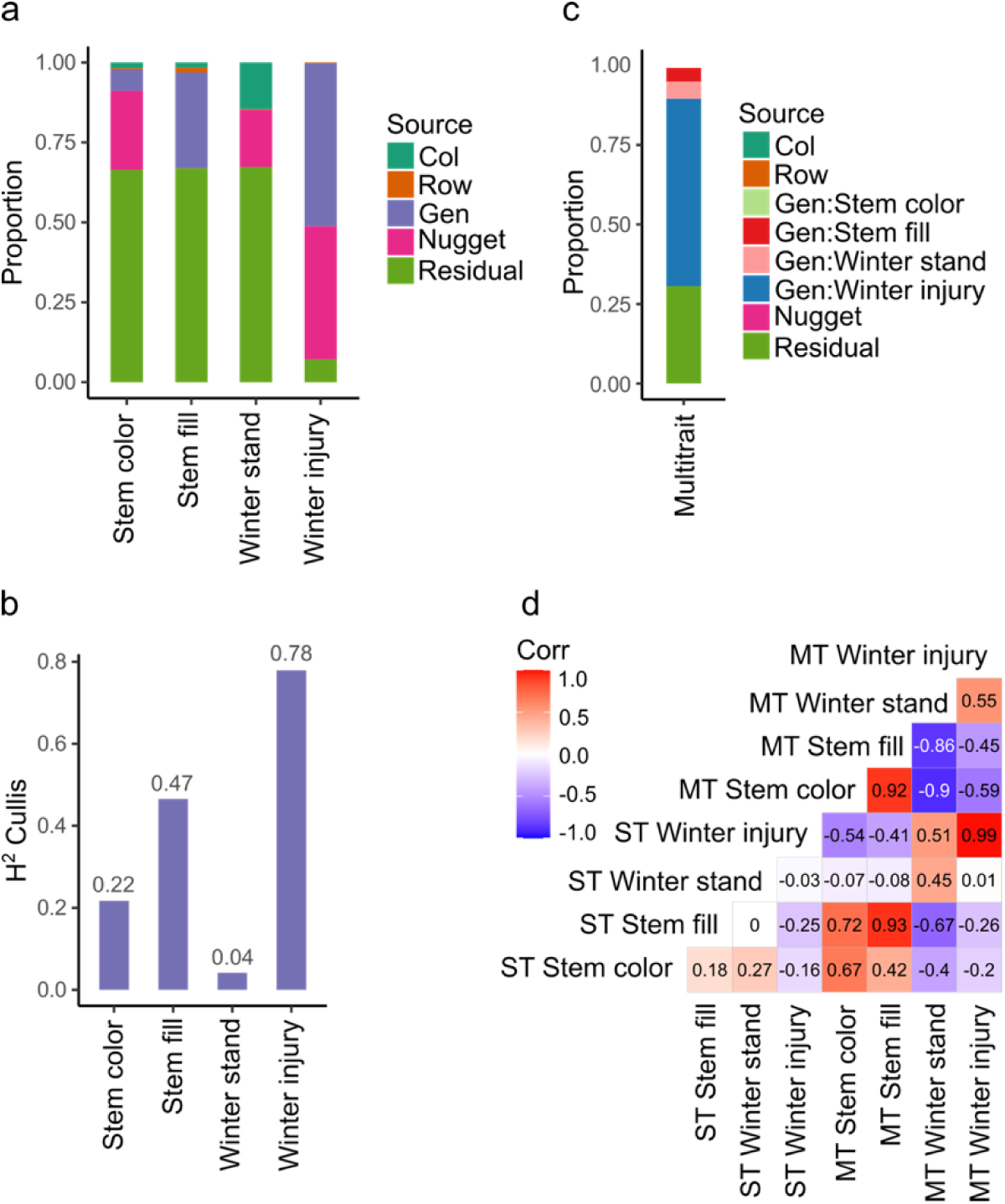
Variance components for random effects in four stem traits. **a.** Proportion of variance components in four traits modeled using a single trait spatial model. Col and Row are spatial variance effect of columns and rows in the field experiment, Gen is genetic variance, Nugget corresponds to the nugget effect, a measure of error variance in spatial models [18]. **b.** Broad sense heritability calculated using the Cullis method [19]. **c.** Proportion of variance components in four traits modeled using multi-trait spatial model. **d.** Pearson correlation of predicted values of traits associated with stem digestibility by univariate (single trait, ST) and multivariate (multi trait, MT) mixed linear modeling. Winter stand, winter standability.

Multivariate or multi-trait (MT) spatial MLM provided different estimates of genetic variance of the traits by fitting them simultaneously in a model. Winter injury kept the highest genetic variance (Vg = 0.42) while stem color has the lowest genetic variance (Vg = 2.88 ×10^-3^), and winter standability increase this value up to 0.04 (Figure 3c). MT-MLM increase the absolute Pearson’s correlation of predicted values between traits. Mean absolute value of Pearson’s correlation in ST-MLM was 0.15 while in MT-MLM was 0.71. BLUEs of winter injury did not change by MT modeling (Pearson’s = 0.99 between ST and MT), but winter standability BLUEs were affected (Pearson’s = 0.45 between ST and MT). The highest Pearson’s correlation in ST modeling was between winter standability and stem color (0.27), and the lowest value was between stem fill and winter injury (-0.25). The highest Pearson’s correlation in MT modeling was between stem fill and stem color (0.92), and the lowest value was between winter standability and stem color (-0.90) (Figure 3d).

### Genome wide association studies

BLUE values from ST and MT spatial modeling were utilized to identify SNPs associated with stem traits. GWAS with ST-BLUEs identified nine significant associated markers (Table 1 and Supplementary Figure 3). Stem color, stem fill, and winter standability each were associated with two markers, while winter injury was associated with three markers. Marker 4_85794609 was associated with two traits: stem color and winter standability. The proportion of explained variance was calculated as R^2^ for each marker; however, all markers exhibited minor phenotypic effects. The highest R^2^ was identified in marker 7_65295546, explaining 6% of winter standability (Table 1). GWAS with MT-BLUEs identified seven significant associated markers. Stem fill and winter standability each were identified with two markers, winter injury with three markers, and no markers were identified for stem color. The mean -log_10_(p-values) of significant markers was higher in MT-BLUEs (5.25) compared with ST-BLUEs (5.16) (Table 1 and Supplementary Figure 3). Twelve out of thirteen markers were in gene coding regions; however, the distribution of the DArTag markers were sparse across the genome and additional candidate genes were annotated in a window of 84 kb according to LD results (Supplementary Table 2).

**Table 1.**
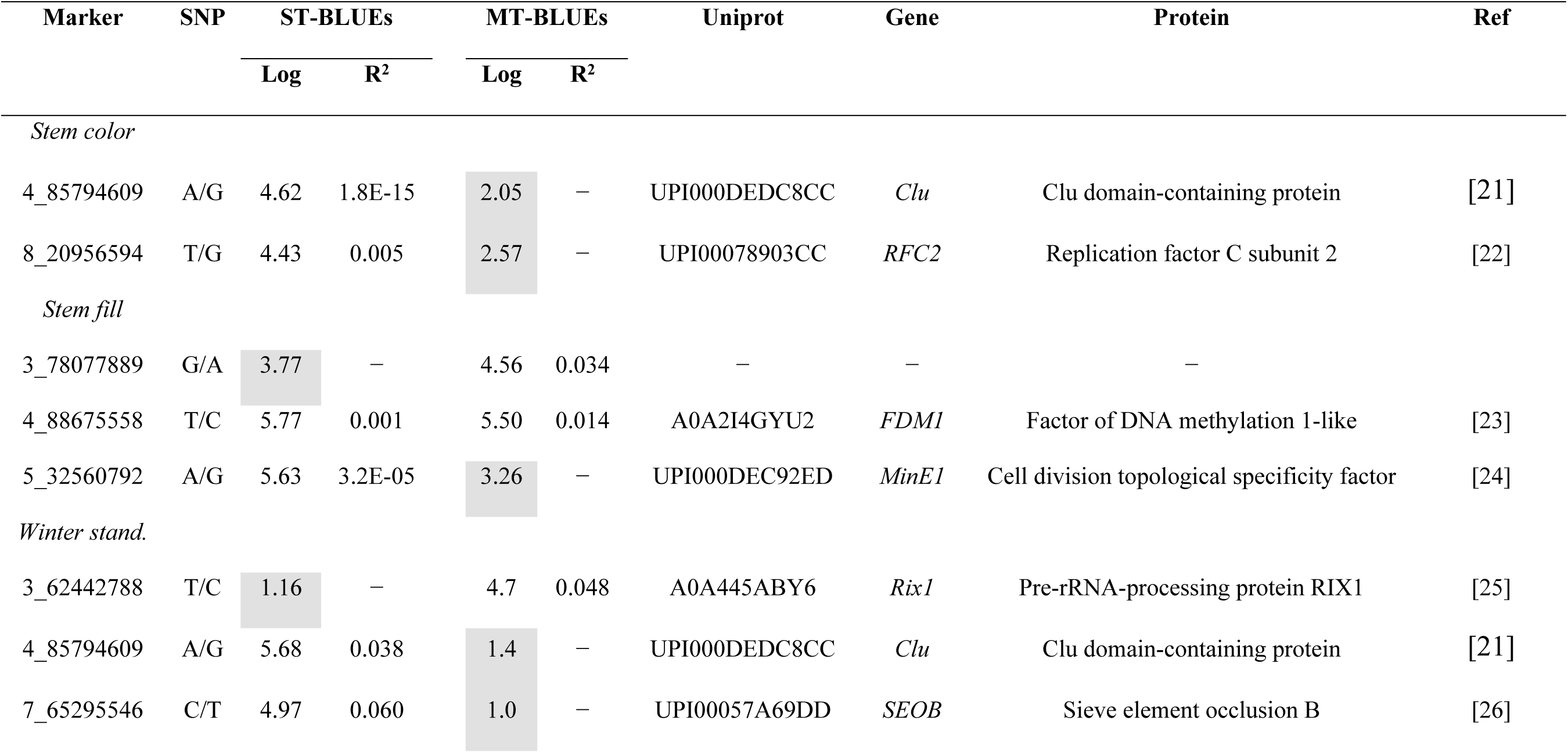
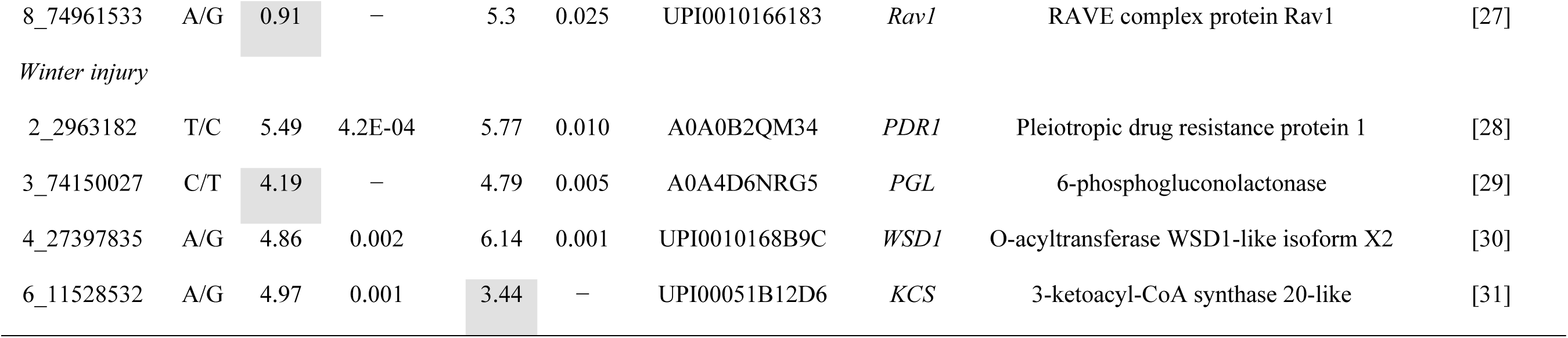
List of significant markers and candidate genes for stem traits. Significant markers were identified using univariate (single trait, ST) and multivariate (multi trait, MT) best linear unbiased estimators (BLUEs). The log is the -log_10_(p-value); R^2^ is the proportion of explained variance. Candidate genes (Gene) were annotated using pan transcriptome information and a monoploid version of *M. sativa* genome [20]. Winter stand. correspond to winter standability. Gray cells indicate markers below the Bonferroni threshold.

Linkage disequilibrium among markers of interest was tested to identify LD blocks in a window of 2 Mb. Six out of nine significant markers had LD blocks with an average size of 393 kb. Four markers (2_2963182, 4_85794609, 4_88675558, and 8_20956594) were in LD blocks of ∼450 kb with other three markers. One marker (4_27397835) was in a block with other two SNPs, and one marker (7_65295546) was in a block with another SNP (Supplementary Figure 4).

### Machine learning predictions

To corroborate the GWAS results, importance scores of markers were calculated using SVM and RF models and compared with GWAS findings. The two markers associated with stem color with ST-BLUEs (4_85794609, 8_20956594) were not associated with MT-BLUEs and had low importance scores (mean of 3.4 and 4.1). Three markers were jointly associated by GWAS using ST and MT-BLUEs in stem fill. Marker 4_88675558 was significantly associated by both ST-BLUEs (5.50) and MT-BLUEs (5.77) and had high importance scores (mean = 81). Marker 3_78077889 was only significantly associated with MT-BLUEs (4.56); however, the mean importance score of SVM and RF was 75.4. Marker 5_32560792 was only significantly associated with ST-BLUEs (5.63), and the mean importance score of SVM and RF was 75.5. Two markers (4_85794609 and 7_65295546) associated with stem fill with MT-BLUEs had the highest importance scores with RF (100 and 99.5). Four markers were significantly associated with winter injury. Markers 2_2963182 and 4_27397835 were significantly associated using both ST-BLUEs (5.49 and 4.86) and MT-BLUEs (5.77 and 6.14) and had high importance scores (mean = 58.9 and 78.9). Two markers (3_74150027, 6_11528532) were significantly associated only with ST-BLUEs or MT-BLUEs, and the mean importance score of SVM and RF was low (mean of 31.1 and 27.2) (Table 2).

**Table 2.** Importance scores of significant markers associated with stem traits. Significant markers were identified through genome-wide association studies (GWAS) using univariate (single trait, ST) and multivariate (multi trait, MT) best linear unbiased estimators (BLUEs). GWAS values correspond to -log_10_(p-values). The importance scores for markers identified by GWAS were calculated using support vector machine (SVM) and random forest (RF) models. Importance scores were scaled from 0 to 100. Winter stand. correspond to winter standability. Gray cells indicate markers below the Bonferroni threshold in GWAS or with a variable importance score <10 in SVM or RF models.

| Marker | Trait | ST-BLUEs |  |  | MT-BLUEs |  |  |
| --- | --- | --- | --- | --- | --- | --- | --- |
|  |  | GWAS | SVM | RF | GWAS | SVM | RF |
| 4_85794609 | Stem color | 4.62 | 1.5 | 5.0 | 2.05 | 4.0 | 3.2 |
| 8_20956594 | Stem color | 4.43 | 1.4 | 8.0 | 2.57 | 5.9 | 1.1 |
| 3_78077889 | Stem fill | 3.77 | 100.0 | 56.8 | 4.56 | 100.0 | 45.0 |
| 4_88675558 | Stem fill | 5.77 | 59.9 | 99.8 | 5.50 | 64.4 | 100.0 |
| 5_32560792 | Stem fill | 5.63 | 78.0 | 100.0 | 3.26 | 80.4 | 43.6 |
| 3_62442788 | Winter stand. | 1.16 | 0.8 | 6.7 | 4.7 | 3.0 | 50.1 |
| 4_85794609 | Winter stand. | 5.68 | 1.5 | 100.0 | 1.4 | 2.4 | 13.4 |
| 7_65295546 | Winter stand. | 4.97 | 82.0 | 99.5 | 1.0 | 0.0 | 18.5 |
| 8_74961533 | Winter stand. | 0.91 | 3.2 | 4.1 | 5.3 | 2.8 | 93.4 |
| 2_2963182 | Winter injury | 5.49 | 34.9 | 75.6 | 5.77 | 39.9 | 85.2 |
| 3_74150027 | Winter injury | 4.19 | 1.9 | 64.2 | 4.79 | 1.8 | 56.6 |
| 4_27397835 | Winter injury | 4.86 | 53.6 | 100.0 | 6.14 | 62.0 | 100.0 |
| 6_11528532 | Winter injury | 4.97 | 13.1 | 39.7 | 3.44 | 13.2 | 42.7 |

Pearson’s correlation of all -log_10_(p-values) from GWAS and all variable importance scores from SVM or RF identified correlated results from multivariate analysis. There are high correlations (>0.8) among importance scores using MT-BLUEs of stem color, stem fill, and winter standability by SVM and GWAS. Winter injury had a high correlation (>0.9) between ST-BLUEs and MT-BLUEs by each model (GWAS, SVM, and RF) but a low correlation among models. For example, the correlation between winter injury SVM and winter injury RF using MT-BLUEs was 0.4 (Supplementary Figure 5a). Plotting variable importance of SVM in the genomic position allowed identification of genomic regions with high importance in stem fill and winter injury. In stem fill, four markers with importance scores >90 were located in a region of 6.4 Mb on chromosome 3 (Supplementary Figure 5b). For winter injury, seven markers with importance scores >50 were located in a region of 12.4 Mb on chromosome 4 (Supplementary Figure 5c).

## Discussion

The alfalfa stem serves as a structural organ supporting all aerial parts and maintaining vascular connections between the roots and leaves. Also, it is an important component of forage yield, representing 50 to 70% of the biomass yield, and influences various traits such as plant height, standability, and digestibility. In this study, we identified 13 molecular markers associated with stem color, stem fill, winter standability, and winter injury in a panel of five divergent stem digestibility populations using a mid-density DArTag platform.

### Population Structure

Previous studies on alfalfa populations and diversity have primarily focused on natural and highly diverse populations from various geographical origins [32,33]. In this study, we utilized principal component analysis (PCA) and structure analysis approaches to visualize the population structure and relationships within germplasm of five populations generated through assortative mating generated over two cycles of bidirectional selection for 16-h and 96-h IVNDFD. PCA and structure analysis revealed clear differentiation between populations with high and low stem digestibility. Structure analysis and interpopulation Rho values identified the highest differences between high and low digestibility populations of C2. Linkage disequilibrium (LD) values at r^2^ = 0.1 ranged from 84 kb to 272 kb, consistent with previous reports in alfalfa [34]. Additionally, LD and Rho values agreed, indicating a greater difference in low digestibility populations (UMN3358 and UMN4019) compared to the C0 population, suggesting a stronger association between alleles in those populations.

### Univariate and Multivariate Modeling

We modeled the morphological traits using spatial mixed linear models employing both univariate (single trait) and multivariate (multi-trait) models. Univariate modeling serves as the initial step due to its simplicity and reduced computational demand in assessing genetic variance and trait heritability. However, univariate models lack the ability to account for potential relationships between traits since they model each trait independently [35]. Therefore, when traits are correlated, multivariate models can provide more accurate predicted values. In our study, we observed low correlations among traits using univariate models (-0.25 to 0.27) and higher correlations in multivariate models (-0.9 to 0.92).

Predicted values of winter injury and stem fill remained consistent across univariate and multivariate modeling (R = 0.99 and 0.93, respectively). However, predicted values of stem color and winter standability differed between the multivariate and univariate models (R = 0.67 and 0.45, respectively), as they were influenced by observations in other traits. These discrepancies affected the molecular markers associated with GWAS. In this study, nine, seven, and three markers were associated with univariate, multivariate, or both modeling approaches, respectively. Multivariate models, designed to capture complex trait relationships, often produce more accurate parameter estimates and predictions than univariate models [36].

### Stem fill

In alfalfa, stem fill was classified into two levels: hollow (0) and solid (1), corresponding to the presence or absence of a hollow in the central pith. In many plants, the death of pith parenchyma cells reduces nutritional value, and palatability; therefore, solid stems increase stem digestibility and nutritive quality. Stem fill can be considered a measure of plant maturity and is related to a reduction in winter standability and resistance to stem lodging in maize [11]. In this study, we identified an inverse correlation between winter standability and solid stems (-0.86). One possible explanation is the maturity stage of alfalfa genotypes. In young stems, highly digestible pith parenchyma cells are intact (score = 1), but there are also lower lignified tissues, decreasing the percentage of erect stems. The unlignified pith parenchyma cells die in mature stems, leading to the formation of a hollow stem (score = 0), and increasing the lignified tissues and the winter standability.

Three markers were associated with stem fill, two of them located in coding regions. The gene containing the marker 4_88675558 encodes a factor of DNA methylation 1-like (*FDM1*). *FDM1* has been reported to control plant and organ size in woodland strawberry by controlling the expression of multiple genes related to the cell cycle and cytoskeleton organization [23]. Marker 5_32560792 was located in a region annotated as a cell division topological specificity factor (*MinE1*). *MinE1* has been reported to control plastid division in *Arabidopsis* [24]. Finally, marker 3_78077889 was not located in a coding region. However, this is one of the four markers with importance scores > 90 with support vector machine (SVM) located in a region of 6.4 Mb on chromosome three (Supplementary Figure 5b). This region contains 301 genes, including 16 transcription factors, such as an ethylene-responsive transcription factor at 73 kb (3_78151631) and a NAC transcription factor at 874 kb (3_77203585). The D gene is a NAC transcription factor and has been reported as responsible for the death of stem pith parenchyma cells in *Sorghum bicolor* [37].

### Stem color and winter standability

Stem color and winter standability were measured at the end of winter or early spring (04/04/22) to relate the effect of winter cold temperatures on stems that grew during the fall. If the stems turned black or brown, it indicated that the tissue was likely killed during the winter. Stem color was categorized as yellow (0) or brown (1) and winter standability was classified from 1 (< 10% of erect stems) to 5 (> 70% erect stems). In this study, we identified a negative correlation between winter standability and stem color (-0.9), i.e., brown stem color was associated with low winter standability. This result could be explained by a loss of water pressure in brown stems, resulting in low standability. Freezing injury in alfalfa results from the pressure exerted by intracellular ice crystals, which disrupt the membrane structure during the thawing process. The damage occurs in older parenchyma cells of the phloem and xylem, as well as in the central pith-like structure of the upper part of the taproot [13]. Multivariate analysis showed significant differences (p-value < 0.05) in C2 populations for stem color and winter standability (Supplementary Figure 2). C2 L×L (UMN4019) had yellow and erect stems while C2 H×H (UMN4016) were brown and prostrate stems, which suggest that low digestible stems can be more tolerant to effect of cold winter temperatures.

Marker 4_85794609 was associated with stem color and winter standability. This marker explains 4% of variation in winter standability and it was located in a locus coding for a Clu-domain-containing protein. Clu proteins are required for mitochondrial subcellular localization [21]. In plant cells, Clu was postulated to control mitochondrial binding and localization by regulating the interaction with microtubules known to assist with cellular structure with water tension [38]. Marker 8_20956594, associated with stem color, was located in a gene annotated as replication factor C subunit 2 (*RFC2*). *RFC2* is involved in DNA replication and repair mechanisms, with high expression in proliferating tissues such as the shoot apical meristem and very weakly in non-proliferating tissues [22]. Marker 7_65295546, which explained 6% of the phenotypic variation in winter standability, was located in a gene annotated as sieve element occlusion B (*SEOB*). *SEO* genes encode P-proteins to facilitate rapid wound sealing after injury, preventing the loss of turgor and photosynthate [26]. In *Medicago truncatula, MtSEO-F1-F3* are specifically expressed in immature sieve elements [39]. Additionally, marker 3_62442788 associated with winter standability was located at 37 kb upstream of other *SEOB* gene (Supplementary Table 2). We can hypothesize that *RFC2* or *SEOB* genes have roles in responding to plant injury and keeping cell turgor for stem standability after cold winter temperatures.

### Winter injury

Winter injury is a consequence of cold winter temperatures. In this work, winter injury was measured in early spring (04/24/23) on a scale from one indicating a dead plant to five indicating no winter injury. Winter injury was the trait with highest Vg and H² in univariate and multivariate analysis, indicating that most of the phenotypic variations of this trait were genetically controlled, giving more confidence in the phenotypic performance to freezing damage during subsequent selection cycles. Although stem color and winter standability were measured during a similar season one year before, their genetic variance and the H² were highest in winter injury. The high H² (0.78) of winter injury indicate that most of the phenotypic variations of this trait were genetically controlled.

Four markers were associated with winter injury, and all of them were in coding regions. Marker 2_2963182 was located in a locus annotated as plasma membrane pleiotropic drug resistance protein 1 (*PDR1*). *PDR1* is an ATP-binding cassette transporter related to plant defense against different fungal and oomycete pathogens [28,40]. Similarly, marker 3_74150027 was located in the pathogenesis-related gene, 6-phosphogluconolactonase *(PGL*). Mutants *pgl3* plants exhibit enhanced resistance to *Pseudomonas syringae* pv. *maculicola* and *Hyaloperonospora arabidopsidis*, and *PGL3* gene is an essential gene for plant size and development [29].

Markers 4_27397835 and 6_11528532 were located in loci annotated as O-acyltransferase WSD1-like isoform X2 (*WSD1*) and 3-ketoacyl-CoA synthase 20-like (*KCS*), respectively. Both genes are involved in the wax biosynthesis pathway [41]. WSD1 functions as wax ester biosynthesis in stems [30], and KCS participates in the synthesis of fatty acid elongation and cuticular wax biosynthetic pathways [31]. Our findings highlight the possible importance of the stem wax biosynthesis pathway in cold tolerance in alfalfa and agree with previous reports. For example, in *Arabidopsis*, the *dewax* mutant showed a greater ability to accumulate waxes under cold acclimation and displayed freezing tolerance at colder temperatures compared with the wild type [42].

### Validation of candidate markers by machine learning models

Breeders need stable targeted markers to track the transmission of traits through different breeding cycles. Here, the stem digestibility panel was genotyped with 2,434 SNP markers. Markers were associated using GWASpoly, which employs the Q+K mixed-effects model incorporating both population structure and co-ancestry among genotypes, as described by Yu et al. (2006) [43]. Although GWAS is a comprehensive approach to systematically search the genome for causal genetic variation, new tools like machine learning (ML) models can enhance the detection of markers associated with traits of interest.

In ML models, all SNPs are ranked based on their variable importance on a scale from 0 to 100. Larger values indicate a greater increase in the prediction error (mean squared error, MSE) when the SNP is randomly permuted, compared to the MSE value prior to permutation. ML has the capability to capture the combined minor effects of multiple genetic markers, allowing for the detection of multi-locus effects during the estimation of the variable importance of SNPs. Additionally, ML models can corroborate GWAS results and enhance the detection of markers associated with traits of interest [17]. In this study, support vector machine (SVM) and random forest (RF) confirmed the significance of 11 out of 13 markers identified by GWAS. Additionally, SVM identified two regions of 6.4 Mb and 12.4 Mb with four and seven markers with high variable importance for stem fill and winter injury, respectively. SVM exhibited lower shrinkage of VI for predictor variables. Therefore, ML enhanced the ability to detect new genetic associations with various traits, addressing the challenges posed by the complex genetic architecture of quantitative traits.

In conclusion, this work demonstrates how the use of multivariate mixed linear models and machine learning can broaden the molecular markers associated with four important traits in alfalfa. The DArTag genotyping platform used in this study has three advantages: 1. It demonstrated that assortative mating during two selection cycles changed the population structure and allele frequency between low and high digestibility populations. 2. Genotypic information was utilized to identify associated markers with four morphological traits using classical Q+K MLM and ML models. 3. The markers identified in this work highlight important mechanisms controlling stem traits such as stem fill or winter survival. DArTag markers identified in this study are highly reproducible in other populations and can facilitate marker-assisted selection to increase winter hardiness or palatability in alfalfa. A next bidirectional selection cycle is in progress and data of IVNDFD such as other forage quality traits will be included in the next part of this project.

## Materials and methods

### Plant materials and field experiment

Plant materials were previously detailed by Xu et al. (2023) [8]. In summary, a parental population (UMN3097) was created by mixing seeds from six commercial alfalfa varieties (5312, Rushmore, Magnagraze, Wintergreen, Windstar, and WL 325HQ). In the selection cycle zero (C0), approximately 2,400 seeds were hand-sown to establish a plant nursery, and the biomass yield from genotypes was harvested at approximately 25% bloom stage. Fresh biomass yield was subsequently dried at 60°C, and stems and leaves were separated. Each ground stem sample underwent scanning via near-infrared spectroscopy to evaluate 16-h and 96-hin vitro neutral detergent fiber digestibility (IVNDFD). The values of 16-h and 96-h IVNDFD were utilized to categorize the UMN3097 population into four groups: 117 plants with high 16-h and 96-h digestibility (H16×H96); 26 plants with low 16-h and low 96-h digestibility (L16×L96); 28 plants with high 16-h and low 96-h digestibility (H16×L96); and 33 plants with low 16-h and high 96-h digestibility (L16×H96).

Plants from C0 were intercrossed by hand tripping flowers without emasculation to generate two populations in selection cycle 1 (C1). Populations UMN3355 and UMN3358 were developed through random intercrossing of the plants H16×H96 and L16×L96, respectively. Seeds from C1 populations were established in a spaced plant nursery, following the procedure outlined for the parental population. Biomass yield was harvested, dried, and separated into leaf and stem fractions, and the stem fraction underwent analysis via near-infrared spectroscopy for 16-h and 96-h IVNDFD. Each of the two C1 populations were categorized into four groups using similar criteria applied in the C0 population.

Values of 16-h and 96-h IVNDFD calculated for the C1 populations were used to develop cycle 2 (C2) populations. UMN4016 was generated by intermating approximately 30 plants of UMN3355 with high 16-h and high 96-h IVNDFD, while UMN4019 was generated by intermating approximately 30 plants of UMN3358 with low 16-h and low 96-h IVNDFD (Supplementary Table 3). One hundred genotypes from each population were established in 2021 at the University of Minnesota Experimental Research Station in Saint Paul, MN. From the original plant, 12 vegetative cuttings were made with three cuttings planted in each replication. The experimental design used was a randomized complete block design with 1,500 plots arranged in three replicates, with 50 rows and 30 columns. To prevent border effects, a border of the alfalfa cultivar Agate was planted around each side of the plots. The plots were managed following best practices, and pesticides were applied as needed.

### Phenotype collection

A set of four stem traits were collected during 2022 and 2023. Stem color and winter standability were collected on 04/04/22 with snow cover from the early spring, stem fill was collected on 07/01/22, and winter injury was collected on 04/24/23. All phenotypic values were collected as categorical traits. Stem color and stem fill had two levels: yellow (0) or brown (1) for stem color and hollow (0) or solid (1) for stem fill. Winter standability was assessed using a modified standability scale ranging from 1 to 5, where 1 represented 0 to 10% erect stems, 2 represented 11 to 30% erect stems, 3 represented 31 to 50% erect stems, 4 represented 51 to 70% erect stems, and 5 represented over 70% erect stems [44]. Winter injury was evaluated on a scale from 1 to 5, where:1 indicated dead plants, 2 indicated less than 15% winter injury, and short plants, 3 indicated less than 10% winter injury, short plants with more branches, 4 indicated less than 5% winter injury, tall plants with many branches, 5 indicated no winter injury, tallest plants with many branches. The injury rate was estimated visually without counting the plants.

### Univariate and multivariate spatial modeling

The phenotypic response of four stem traits were modeled separately by single trait (ST, i.e. univariate) modeling and conjointly by multi trait (MT, i.e. multivariate) modeling. Each phenotypic response consisted of 1,500 observations corresponding to the plots indexed by 50 rows and 30 columns in a contiguous field array, with three complete blocks and 500 genotypes. Four traits were modeled separately by a univariate mixed model y = **X**β +**Z**u + ε, where y is the response variable, **X** and **Z** were incidence matrices for fixed and random effects respectively, β was the vector of fixed effects associated to genotypes, u was the vector of random effects associated to genotypes, rows and columns, and ε was the vector of residuals. It is assumed that the random terms (u,ε) are pairwise independent with zero mean and variance-covariance matrices.

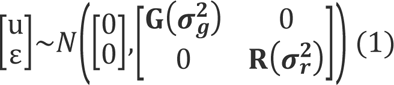

where **G** and **R** are covariance matrices of vectors of genetic 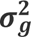 and residual 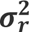 variance, respectively. The form of the covariance matrix for the plot errors (**R**) is given by a separable first-order autoregressive model using the field row and column positions [45]. The nugget effect was included to measure the variance to-error variance ratio.

Multivariate spatial modeling uses the same parameters described in the univariate modeling, but data were ordered by traits (*t*) within units in a variance matrix **I**_*n*_**⨂Σ**, where **Σ**^(*t*×*t*)^was a factor analytic variance matrix. The error structure was specified with independent units and an unstructured variance matrix.

The ASReml-R software was utilized to estimate the variance parameters, best linear unbiased estimates (BLUEs) of the fixed effects, and empirical best linear unbiased predictors of the random effects [18]. Broad sense heritability was calculated using the Cullis method [19] assuming the genetic term as random effect:

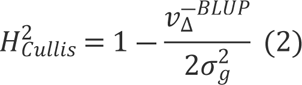

where 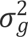 is the genetic variance and 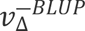 is the average standard error of the genotypic BLUPs calculated from ASReml-R spatial model.

### DArTag genotyping

Genomic representations of 1,502 genotypes were generated by collecting two to three leaflets (∼100 mg) per genotype for DNA extraction and genotyping. Leaf tissues were sent to Intertek (Intertek; Alnarp, Sweden) for DNA extraction [46]. The DNA samples were then sent to Diversity Arrays Technology Ltd. (DArT; Canberra, Australia) [47] for genotyping using the 3K DArTag genotyping platform developed by Breeding Insight (Breeding Insight; Ithaca, NY, USA), Cornell University [14]. The targeted regions were amplified by PCR with sample-specific barcodes and then sequenced to identify the present polymorphisms.

DArTag generated the Allele Match Count Collapsed (AMCC) file containing the read depths of targeted SNPs in the alfalfa SNP array. The AMCC file was then converted into a RADdata object using the ‘readDArTag’ function to export discrete genotypes in a molecular matrix with the polyRAD R package v.2.0.0 [48]. SNPs were filtered based on minor allele frequency (MAF) > 0.05, and the number of redundant markers with identical genotype calls were reduced using the ‘snp.pruning’ function of the ASRgenomics v1.1.4 R package [49]. This process resulted in a genotypic matrix with 2,434 SNPs×1,502 genotypes.

### Population structure

Principal Component Analysis (PCA) was conducted using the built-in R function ‘prcomp’ with the setting ‘scale = TRU’. For population structure analysis, Structure v.2.3.4 software was utilized [50]. To determine the appropriate number of inferred clusters to model the data, ten independent runs were conducted for each number of subpopulations (K) ranging from 2 to 10. The burn-in length was set to 10,000, and the Markov Chain Monte Carlo (MCMC) length was set to 10,000 as well. To identify the optimal number of clusters (K), the Evanno method was employed, where the result with the largest LnP(D) and the smallest K values is considered optimal [51]. The optimal K value and the population structure bar plot at K = 2 were generated using the pophelper R package v.2.3.1 [52]. The cluster values obtained from Structure were included as covariates in the genome-wide association study (GWAS).

To evaluate genetic diversity, observed heterozygosity (H_O_), expected heterozygosity within a population (H_S_), and interpopulation differentiation measured with the Rho parameter were calculated using the software GenoDive v.3.0 [53].

### Linkage disequilibrium

Linkage disequilibrium (LD) between each pair of SNPs was estimated using squared allele-frequency correlations (*r*^2^) calculated with the ldsep R package v.2.1.5 [54]. The rate of LD decay was estimated using *r*^2^and the distances in base pairs (bp) based on the *M. sativa* cultivar XinJiangDaYe reference genome [55] using a nonlinear model [56]. The expected *r*^2^value was *E*(*r*^2^) = 1/(1 + *C*), where *C* = 4*ad*, where *a* is an estimated regression coefficient and *d* is the physical distance in bp. Assuming a low level of mutation and finite sample size *n*, the expectation becomes [57]:

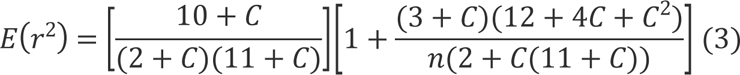

To evaluate the consistency of LD, global LD decay and estimated LD values for each chromosome were defined as the distance at *r*^2^ = 0.1.

### Genome-wide association study and functional annotation

GWAS was conducted using GWASpoly v2.13 [58] using the Q+K mixed linear model, which conducts single locus analysis incorporating structure information (Q) and a kinship matrix (K) in the model to reduce false positives resulting from population structure and family relatedness [59]. The cluster values obtained from Structure were used as Q, while K was calculated by GWASpoly. The -log_10_(p-values) were corrected by Bonferroni method at a cutoff value of 5% to identify SNPs significantly associated with the traits. Subsequently, significantly associated markers were annotated against the UniProt100 database [60] using the reference transcriptome dataset for *M. sativa* [61] in a window of 84 kb according to LD results.

### Machine learning models

Support vector machine (SVM) and random forest (RF) models were implemented to identify linear and non-linear marker-trait associations by fitting all markers simultaneously. Variable importance scores of SNPs were calculated using to understand how the SNPs in the testing model contribute to the prediction model. Variable importance scores were compared with -log_10_(p-values) from GWAS.

SVM find the best hyperplane with the maximal margin in an *p*−dimensional space with respect to a given response variable [62]. A hyperplane refers to a straight line in a high-dimensional or *p*-dimensional space, such as the genotypic matrix with *p* SNPs, where the response variables are the *n* BLUEs. In SVM, each *n*-dimensional input vector (**x**_*i*_) of *p* SNP markers is associated with a y_*i*_phenotypic response where **x**_*i*_ ∈ ℝ^*p*^ and y_*i*_ ∈ ℝ. The following linear regression is performed *f*(**x**) = 〈*w*,*x*〉 +*b*, where 〈*w*,*x*〉 is the inner product between vectors *w*,*x* ∈ ℝ^*p*^, *w* is the slope and *b* is the intercept of the hyperplane to be estimated [63].

RF independently and uniformly samples with replacement the training data (*L*) to create a bootstrap dataset (*L\**) from which a decision tree is grown. The process is repeated *K* replicates to produce *K* decision trees which form a RF. In each decision tree, multiple binary filters are applied to create a branching structure, forming a tree-like hierarchy. Each point where samples are partitioned is termed a decision node. To estimate the importance scores of SNPs, RF calculates the Gini importance which quantifies the difference between a node’s impurity and the weighted sum of the impurities of the two descendent nodes. The Gini variable importance (*VI*) is given by

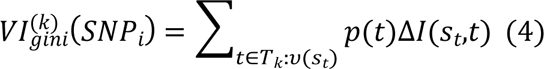

where *T*_*k*_is the number of nodes in the *k^th^* tree, *υ*(*S*_*t*_) is the variable used in the split *S*_*t*_, *p*(*t*) is the fraction of the samples reaching the node *t*, and Δ*I* is the decrease in impurity for all the nodes *t* where the *SNP_i_* is split (*S*_*t*_). SVM and RF models were performed using a ten-fold cross-validation with the caret R package v6.0.94 [64] and the SNP’s importance scores were ranked from 0 to 100 according to the ranks of each SNP’s impact in the trait.

## Acknowledgments

We acknowledge Joshua Larson, Melinda Dornbusch, Keith Henjum, and Ted Jeo for technical assistance in collecting the phenotypic data and maintaining the populations.

## Authors contributions

C.A.M.: Manuscript preparation and data analysis; D.H.J.: Population generation, conceived and outlined this research, and review and editing; D.Z.: DArTag genotyping and review and editing; M.L.: DArTag genotyping and review and editing; C.T.B.: DArTag genotyping and review and editing; M.J.S.: DArTag genotyping and review and editing; D.S.: Population generation, conceived and outlined this research, and review and editing; Z.Y.: Population generation, conceived and outlined this research, and manuscript preparation. All authors have read and agreed to submit the manuscript.

## Data availability statement

Large data sets, including genotypic matrix, BLUP values, Structure cluster membership values, and variable importance scores from support vector machine and random forest are available in figshare (https://doi.org/10.6084/m9.figshare.25686405.v1).

## Disclaimer

Mention of trade names or commercial products in this publication is solely for the purpose of providing specific information and does not imply recommendation or endorsement by the U.S. Department of Agriculture. USDA is an equal opportunity provider and employer.

## Financial Disclosure Statement

This research was supported in part by the U.S. Department of Agriculture, Agricultural Research Service. Breeding Insight was funded by U.S. Department of Agriculture, under agreement numbers (8062-21000-043-004-A, 8062-21000-052-002-A, and 8062-21000-052-003-A).

## Competing interests

The authors have declared that no competing interests exist.

## Notes

### Competing Interest Statement

The authors have declared no competing interest.

